# Sex chromosome dosage compensation in a sex reversing skink is not influenced by sexual phenotype

**DOI:** 10.1101/2023.08.24.554710

**Authors:** Benjamin J. Hanrahan, J King Chang, Ashley M. Milton, Nicholas C. Lister, Duminda S.B. Dissanayake, Jillian M. Hammond, Andre L.M. Reis, Ira W. Deveson, Aurora Ruiz-Herrera, Hardip R. Patel, Jennifer A. Marshall Graves, Arthur Georges, Paul D. Waters

## Abstract

**Background:** Lizards have sex determination systems that can differ between even closely related species. These include XY and ZW systems, and thermolabile systems where genes and temperature interact to determine sex. The eastern three-lined skink (*Bassiana duperreyi*) has a differentiated XY sex determination system, in which low temperature incubation during development can cause female to male sex reversal, producing XX males. This provides a unique opportunity to investigate how genotype and sexual phenotype affect dosage compensation.

**Results:** Here, we present a draft genome assembly of the Eastern three-lined skink generated from nanopore sequencing. We also generated transcriptomes from brain and heart tissue of normal adult males and females, along with brain tissue of sex-reversed XX males. We observed partial dosage compensation between XX females and XY males in both brain and heart, with median gene expression from the X in normal males being 0.7 times that of normal females. In brain of sex reversed XX males the median X chromosome output matched that of the normal XX female level, and not that of normal XY males.

**Conclusions:** Partial dosage compensation in the Eastern three-lined skink is similar to several other species of lizard. However, here for the first time we describe dosage compensation in a lizard with natural sex reversal, and show that in sex reversed individuals dosage compensation of the X chromosome follows genotypic sex and not phenotypic sex.

## INTRODUCTION

Sex chromosome dosage compensation in mammals was proposed to have evolved in response to loss of gene function from the mammalian Y chromosome. Gene expression from the single X in males should need to be upregulated to restore ancestral autosomal levels. This X upregulation carried through to females resulting in disproportionately high X expression in XX females (1), and this was countered by X chromosome inactivation (XCI) to silence one X chromosome in the somatic cells of females (2). This classic model of dosage compensation appears to hold true for marsupial mammals, but transcriptional upregulation of the single X appears incomplete in eutherian mammals (3, 4). In monotremes, median gene expression from the X chromosome is increased compared to the autosomes in XY males, as it is for the bird Z in ZW females. However, global X (or Z) transcriptional output is not evenly balanced between the sexes (5).

Mammals and birds have ancient and relatively stable sex chromosomes and sex determination systems. Most mammals have a conserved XY system and birds have a conserved ZW system. In contrast, reptiles display a wide array of sex determination and sex chromosome systems, even in closely related species (6, 7). These organisms exhibit not only XY and ZW genetic sex determination (GSD) systems, but also temperature-dependent sex determination (TSD), whereby the sex of offspring is determined by the temperature at which the egg is incubated during a thermosensitive window. GSD and TSD systems are often observed in closely related lizard species, and some species have a GSD system that can be overridden by temperature to cause sex reversal (8–11).

Studies of reptile dosage compensation have revealed a variety of non-canonical dosage compensation systems. In snakes, there are reports of partial dosage compensation of the Z chromosome by upregulation of Z-borne genes in females (12, 13). The green anole (*Anolis carolinensis*), a lizard with an XY sex chromosome system, has complete dosage compensation via 2-fold upregulation of the single X in males to match expression with the ancestral autosomal level and balancing expression with the two Xs in females (14), which is mediated by a lncRNA (15). The blacktail brush lizard also has two-fold upregulation of the single X in males (16), but with unknown mechanism. In contrast, the Komodo dragon (*Varanus komodoensis*), which has one of the oldest ZW sex chromosome systems, shows no evidence of dosage compensation of Z-specific genes (17). Partial dosage compensation has been observed in the brown basilisk (*Basiliscus vittatus*) such that the XY males had expression intermediate between full dosage compensation of the two X’s in females, and half the expression (18). Incomplete dosage compensation was also observed in Gila monster (19), which has a ZW sex chromosome system.

The scincid lizard *Bassiana duperreyi* (Australian eastern three-lined skink) has a differentiated XY sex chromosome system (20) that is thought to predate skink radiation (21). However, the genetic sex determining switch can be overridden by temperature (8, 9, 11). Reduced incubation temperature of eggs during the thermosensitive period results in sex reversed XX males in the adult population and the natural nests of both captive and wild populations (22, 23). This gives rise to three different genotypic and phenotypic sexes: XX females, XY males and sex reversed XX males. Therefore, *B. duperreyi* provides a unique opportunity not only to examine dosage compensation in a reptile with differentiated sex chromosomes, but to also test if dosage compensation in sex reversed XX males follows genotype or sexual phenotype.

Here we present a chromosome level *B. duperreyi* draft genome assembly from a male individual, in which we have identified both the X and Y chromosomes. We generated mRNA sequence data from brain and heart tissue of genotypically normal XX females and XY males, and brain tissue of XX male individuals to examine the effect of genotype and phenotype on dosage compensation. In brain and heart of normal XY male and XX female individuals, there was partial dosage compensation, such that the single X of XY males was overexpressed. Sex reversed XX males were observed to have the same median X gene expression as XX females, showing for the first time that dosage of the X follows genotype in a sex reversed lizard.

## RESULTS

### Genome assembly summary

Genome length was 1.485 Gbp, assembled into 5,369 scaffolds. The six largest scaffolds (288.7 - 75.5 Mbp) corresponded to autosome macrochromosomes (from 1 to 6), with the next six largest scaffolds (54.7 – 18.1 Mbp) representing the microchromosomes (from 7 to 14) (Figure 1A, Table 1). Based on HiC contacts, contig 235 was found to contain merged microchromosomes, likely chromosomes 12, 13 and 14 (Figure 1A, inset). Including the sex chromosome scaffolds this comprised all the chromosomes in *B. duperreyi* (2n = 30). The chromosome-scale scaffolds comprised 98.5% of the total assembled sequence. BUSCO analysis showed 96.0% complete conserved orthologs in the tetrapod database and 96.6% complete conserved orthologs in the vertebrate database (Table 2).

**Figure 1.**
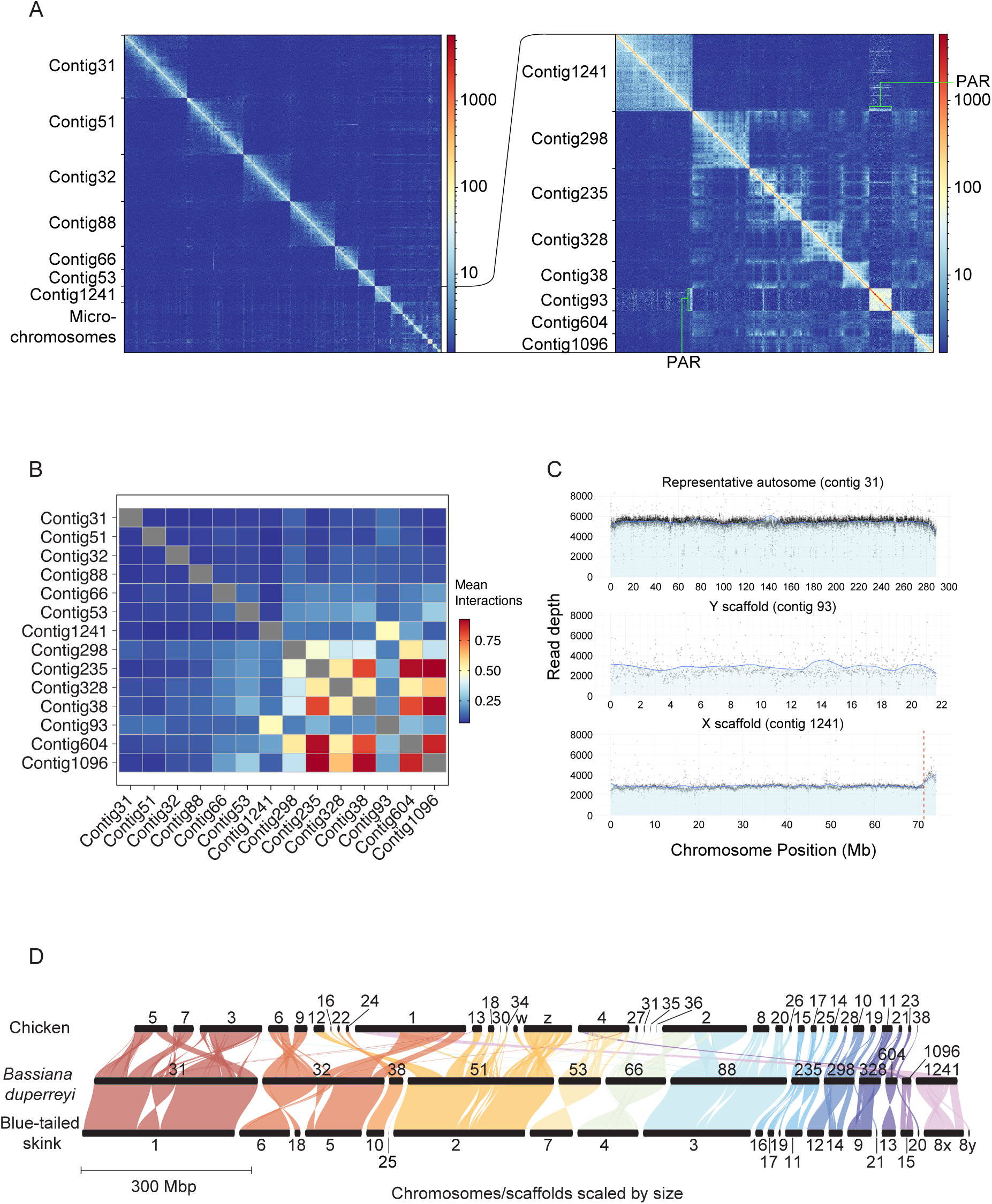
Genome summary of the *Bassiana duperreyi* assembly. **A)** Hi-C contact map generated with HiC explorer (53) showing scaffold (chromosome) boundaries. Inset: magnified view of the HiC contact map for microchromosomes with size 18.1 – 55.4 Mb, and the sex chromosomes. Number of contacts are between 100 kbp bins. Below: mean inter-chromosomal interactions for 100 kbp windows between each chromosome. **B)** Number of reads mapped in 20kb windows of male Illumina genome sequencing on a representative autosome (contig 31), the putative Y (contig 93) and X contigs (contig 1241). Vertical red line is the PAR boundary on the X contig. **C)** Synteny across chicken, bassiana and blue tailed skink (*Cryptoblepharus egeriae*) genomes, generated with GENESPACE (54). The bassiana Y scaffold (contig 93, not shown) shares no synteny with the other two species.

**Table 1.**
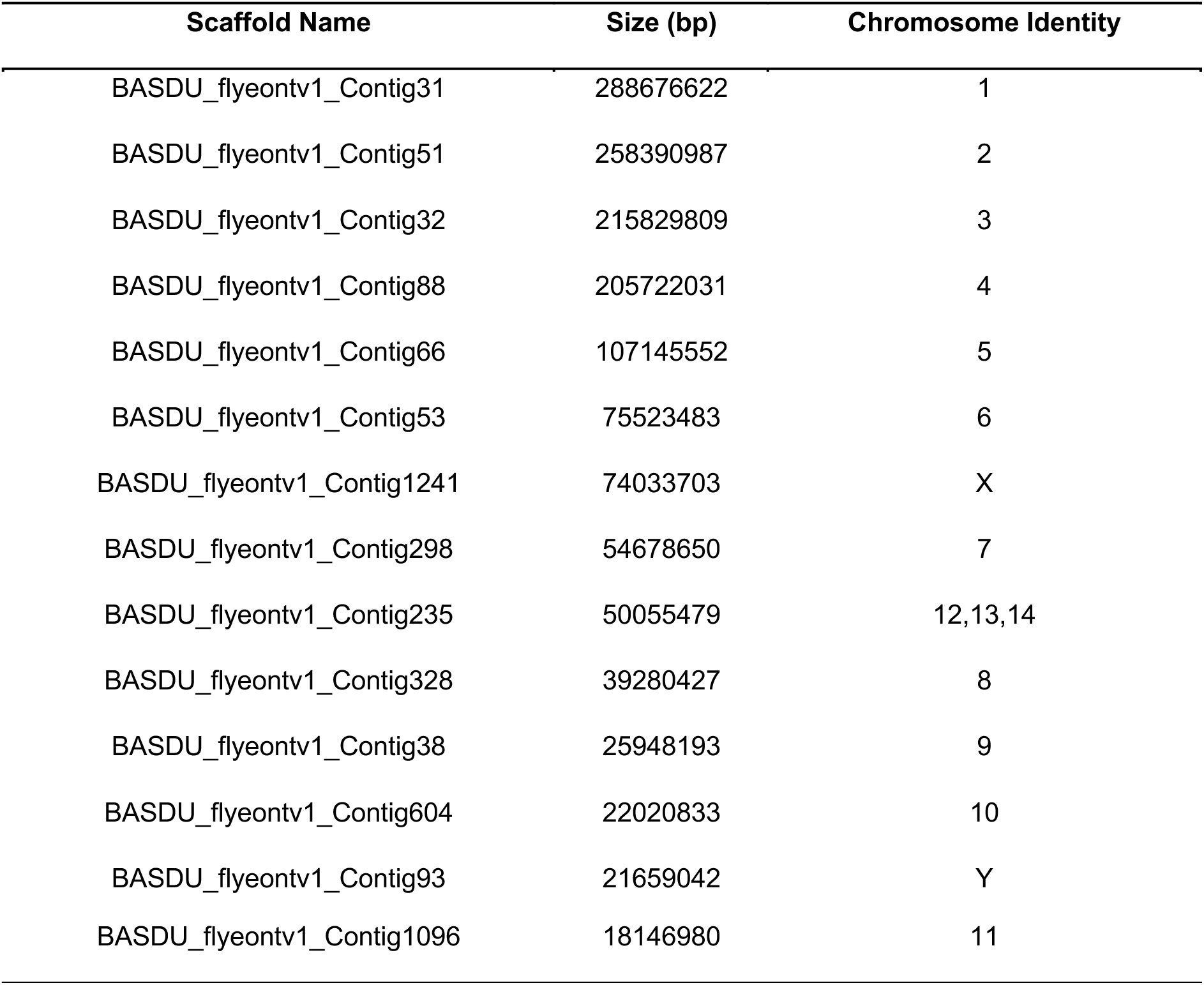
Chromosome identity based on scaffold size.

**Table 2.** BUSCO assessment of the *B. duperreyi* genome.

| BUSCO<br>Assessment | Total | Complete<br>and single-<br>copy | Complete<br>and<br>duplicated | Fragmented | Missing | Groups<br>searched |
| --- | --- | --- | --- | --- | --- | --- |
| *tetrapod_odb10* | 5156 | 5098<br>(96.0%) | 58 (1.1%) | 44 (0.8%) | 110<br>(2.1%) | 38 |
| *vertebrate_odb10* | 3290 | 3240<br>(96.6%) | 50 (1.5%) | 32 (1.0%) | 32<br>(0.9%) | 67 |

### Sex chromosome summary

Contig 93 was identified as the Y chromosome by BLASTn with male specific *B. duperreyi* k-mer contigs (Supplemental Figure 1). Read depth analysis of short read Illumina whole genome sequencing from a male individual confirmed contig 1241 as the X (Figure 1C), and revealed the pseudoautosomal region (PAR) boundary location at which read coverage increased compared to the X and Y specific regions. The X and Y chromosomes were 71.2 Mbp and 21.7 Mbp, respectively, and shared a small PAR of approximately 2.8 Mbp that was assembled onto the X scaffold (Figure 1C). The Y chromosome had more interactions with the macrochromosomes, than the X chromosome (Figure 1B). Fewer interactions of the X with autosomes is similar to eutherian XY systems that also have fewer interactions between the X chromosome and autosomes (24). As expected, microchromosomes interacted with each other more than with the macrochromosomes (25).

Annotation identified a total of 13,979 genes in the genome. Of these, 667 were X-specific genes, sharing homology with chicken chromosome 1 (Figure 1D). A total of 62 genes were annotated on the male specific region of the Y: 47 of these were X/Y shared, with the remaining 15 having no detected X-borne partner. A total of 39 genes were annotated in the PAR.

There were no XY shared or Y specific genes with a known role in the sex determining pathway of other vertebrates (Supplemental Table 1). The only gene conserved on the Y chromosome of the Christmas Island blue-tailed skink was *Ifit5*, which encodes for an interferon that has not been implicated in sex determining pathways. Gene Onology (GO) analysis of the 15 Y specific genes showed no enrichment of genes involved in male related functions. In fact, these genes were not enriched for any biological function. GO analysis of XY shared genes, as well as all genes on the Y (XY shared and Y specific genes), revealed an enrichment of genes involved in histone H4 acetyltransferase activity (Supplemental Tables 2 and 3). In summary, there was no candidate sex determining gene identified.

The karyotypic difference between the X and Y (20), and the disparity between their gene contents, confirmed their differentiation. This suggested an unbalanced output of X-specific genes in XX females and XY males that might be balanced by a sex chromosome dosage compensation mechanism.

### Dosage compensation of the X chromosome

In brain and heart, median gene expression from the autosome scaffolds and PAR was equivalent between XY males and XX females (Figure 2A). However, median X chromosome transcriptional output in XY males was reduced compared to XX females (to 70% in brain and 65% in heart: Mood’s median test, p < 0.0001).

**Figure 2.**
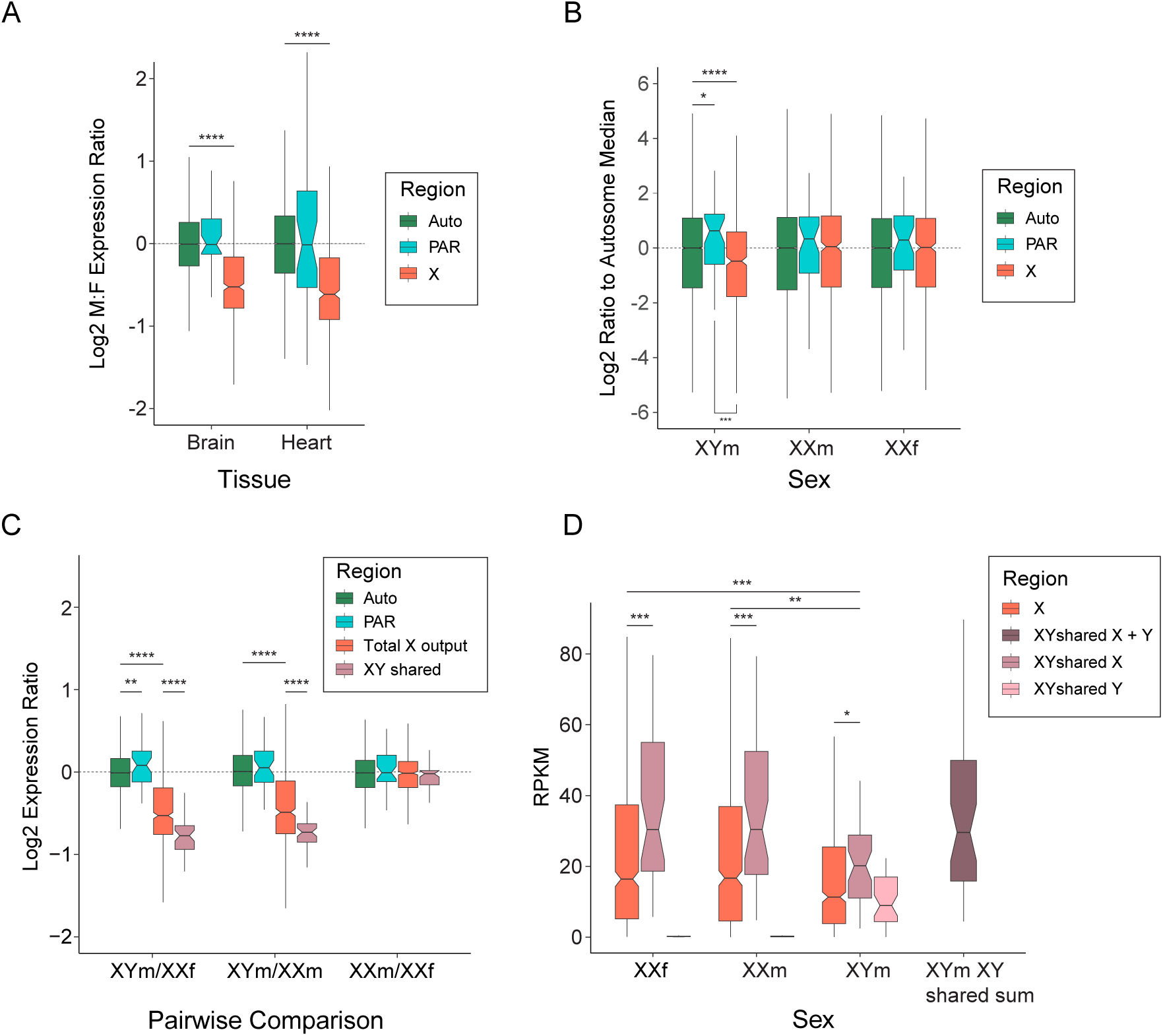
Dosage of gene expression between sexes. **A)** Median expression male (XY) to female (XX) ratios were calculated for adult brain and heart tissue (n=1 for both sexes and tissues). A ratio above zero indicates higher expression in male and a ratio below zero indicates higher expression in female. Median is plotted in the box with exact value above each median. Boxes represent the middle 50% of the data, and whiskers represent 1.5 times the interquartile range. Outliers not plotted. **B)** Brain transcriptomes were used to calculate mean RPKM for each sex condition (XYm, n=3; XXm, n=2; and XXf, n=3) and plotted for autosomal, X specific and PAR genes as a ratio to autosomal median. Ratio to the autosomal median is plotted in the box with exact value above each median. Permutation tests were used to calculate whether median PAR and X specific RPKMs were statistically different from the median autosome RPKM, as well as if the median PAR RPKM was different from the median X specific RPKM (**** p<0.0001, *** p<0.001, * p<0.05). **C)** Median expression ratios were calculated for pairwise comparisons of the three sexes (XYm, XXm and XXf). Genomic regions were separated and plotted as follows; autosomes (green), PAR (blue), total X output (red) and XY shared genes (pink). Mood’s median tests were used to calculate if the ratios for the autosomes and X, as well as Xs between sexes, were statistically different (**** p<0.0001). **D)** As above mean RPKM values were calculated for each sex conditions and represented as a boxplot. Genomic regions on the X chromosome were separated as follows; X (red), XY shared Y (light pink), XY shared X (pink) and sum of X and Y ametologues (dark pink).

We next tested if phenotypic sex influenced total transcriptional output from the X versus the autosomes in brain. We measured the ratios of median X specific to autosome gene expression levels within the three genotypic and phenotypic sexes. In XX females and XX males there was no significant difference between X and autosomal expression levels. In XY males, median transcriptional output from the X was 72% that of the autosomal median (permutation test, p < 0.0001) (Figure 2B).

We then compared transcriptional output of the X specific region between the three phenotypic and genotypic sexes. Median gene expression from autosome scaffolds of XX females, XY males and sex reversed XX males were equivalent (Figure 2C), individual autosome scaffolds were also equivalent (Supplemental Figure 2). PAR region showed slightly higher expression in XY males, however this was only significant in the XYm/XXf comparison. As observed for heart and brain above, median gene expression from the X chromosome was significantly lower in normal XY males compared to XX females (down to 69%: Mood’s median test, p < 0.0001) (Figure 2C). Median gene expression from the X chromosome was also lower in XY males than XX males (down to 71%: Mood’s median test, p < 0.0001). These ratios are both greater than the 50% expected from a non-dosage compensated X chromosome, implying significant upregulation of the single X in XY males relative to both XX females and XX males (Figure 2C). This upregulation of genes on the X in XY males compared to both XX females and XX males was not region specific on the X (Supplemental Figure 3), nor was it restricted to a subset of genes (Supplemental Figure 4).

We also compared expression from the X between sex reversed XX males and normal XX females. Both have two X chromosomes, so we could directly address if sex chromosome dosage compensation strictly follows genotype, or was also influenced by phenotype. We observed that XX males and XX females had equivalent transcriptional output from the X, with the same expression ratio as the autosomes (Mood’s median test, p = 0.886) (Figure 2C), and that the median output from the X was not different compared to autosomes in either XX males (permutation test, p = 0.333) or XX females (permutation test, p = 0.433) (Figure 2B).

Having established that sex chromosome dosage compensation followed genotype, we investigated expression of X genes with a Y gametologue. These X genes (n=42) had significantly lower total transcriptional output in XY males than did XX females (Mood’s median test, p < 0.0001) and XX males (Mood’s median test, p < 0.0001) (Figure 2C). However, when counts for X and Y gametologues in XY males were summed their expression was equal to that of X gametologues in XX males and XX females (Figure 2D). Median expression of the XY shared genes was found to be higher than total X gene expression in all sexes (Mood’s median test: XXf, p < 0.001; XXm, p < 0.001; XYm, p < 0.05) (Figure 2D).

## DISCUSSION

Here we present a draft assembly for *B. duperreyi* with chromosome level scaffolds that included both X and Y chromosomes. Gene expression of the X chromosome between XY males and XX females revealed partial dosage compensation of the X in XY males. Dosage of the genes on the X chromosome in sex reversed XX males was equal to the X in XX females, therefore sex dosage compensation followed genotype.

In normal XY male *B. duperreyi* there was partial dosage compensation of the X chromosome (Figure 2C), with median X chromosome gene expression 69% that of XX females. Therefore, the single X chromosome is being over-expressed by a factor of approximately 1.45. This is similar to observations in non-therian mammal vertebrates such as platypus (5). It is also similar to the dosage system seen in the brown basilisk (18) and Gila monster (19). However, it is different from dosage compensation in other lizard species; the absence of sex chromosome dosage compensation in the ZW system of the Komodo dragon (17), and complete dosage compensation in the green anole (14) and the blacktail brush lizard (16).

Thus, we demonstrate that *B. duperreyi* represents a partial dosage compensation system in a lizard that is chromosome-wide with no region-specific effect. Thus different dosage compensation systems have evolved repeatedly and independently in different lizard lineages, perhaps depending on the dosage sensitivity of the progenitor autosomes from which the sex chromosome systems evolved. Study of more lizards will elucidate the breadth of reptile sex chromosome dosage compensation and highlight different evolutionary strategies.

As well as comparing normal XY males and XX females, for the first time, we describe dosage compensation in a model with naturally occurring sex reversal. We observed that gene expression in the sex reversed XX males was equivalent to that in normal XX females, with median X expression 99% that of XX females and 140% that of XY males. Therefore, it is genotype, as opposed to sexual phenotype, that dictates sex chromosome dosage compensation in *B. duperreyi*. This is the first example showing dosage compensation in a sex reversed animal matches that of the sex with the same genotype. It remains to be seen if the offspring of normal XX females and sex reversed XX males also follow this trend of dosage compensation matching normal females, as it is unknown if sex reversed XX males are reproductively viable (22, 26), though it is likely this is the case.

Temperature-induced sex reversal in lizards is not unique to *B. duperreyi,* having been observed in several species (8, 10). It is possible that sex reversal in lizards is more widespread than previously believed, as it has been systematically investigated in only a few species (27), and there may be numerous other species whose sex reversal status is yet to be confirmed (see 28, 29, 30). Examining dosage compensation in other sex reversing models will confirm whether our finding that phenotype does not influence sex chromosome dosage is ubiquitous.

Recent evidence suggests that in systems with partial or incomplete dosage compensation in the transcriptome, the abundance of proteins can remain balanced (31). This should be investigated further in organisms such as *B. duperreyi,* brown basilisk, Gila monster and komodo dragon that show partial or lack of dosage compensation in the transcriptome, to observe if the sex chromosome output is balanced in the proteome.

## CONCLUSIONS

Here we analysed transcriptomes from brain and heart of adult *B. duperreyi* to show that in canonical XY males there is partial dosage compensation of the X chromosome. The brain transcriptome of hatchling brains also revealed that sex reversed XX males have the same transcriptomic output from the X chromosome as canonical XX females. This is the first description of X chromosome dosage in a naturally sex-reversing system and demonstrates that, in *B. duperreyi*, dosage follows the underlying genotype, regardless of an individual’s sex.

## MATERIALS AND METHODS

### Sample collection and sexing

In December 2020, samples (eggs) of alpine *B. duperreyi* were collected from nests in the field location within the Brindabella Range (Piccadilly Circus – 1240 m a.s.l., 35°21’42.0“S 148°48’12.5”E) after ensuring that approximately 90% of the development period had passed in natural conditions (9). The eggs were then collected and transported to the University of Canberra, where they were buried in moist vermiculite with a ratio of 4 parts water to 5 parts vermiculite by weight, and placed in incubators (LabWit, ZXSDR1090) that maintained 23°C, which produces a balanced sex ratio (9). Details egg collection methods (23) and a description of the alpine study site can be found in (32).

To determine phenotypic sex of the hatchlings, tail bases of 7-day old hatchlings were squeezed to evert the hemipenes (33) and sex was checked again by hemipene transillumination after 5 weeks (23). Tail snips were collected to determine the genotype of the lizards. Genotypic sex was determined for *B. duperreyi* using polymerase chain reaction (PCR)-based molecular sex tests from extracted DNA collected from tissue samples. DNA purity was determined using a NanoDrop 1000 spectrophotometer (NanoDrop Technologies Inc., Wilmington, DE, USA) and quantified using the Qubit 2.0 Fluorometric Quantitation (Invitrogen, Life technologies, Sydney, N.S.W., Australia). The sex-reversal status was determined for *B. duperreyi* using PCR as described by Dissanayake et al. (20), where the genotypic sex was identified based on Y-specific markers allowing identification of XY males (XYm) and XX males (XXm). The lizards were euthanised via intraperitoneal injection of sodium pentobarbitone (100–150 μg/g body weight) as approved by the University of Canberra Animal Ethics Committee (AEC 17–26).

### RNA extraction and sequencing

Total RNA was extracted from the brain tissue of three XY males (XYm), three XX females (XXfm), and two XX males (XXm). Tissue extracts were homogenized using T10 Basic ULTRA-TURRAX® Homogenizer (IKA, Staufen im Breisgau, Germany) and extracted using TRIzol reagent following the manufacturer’s instructions, purifying with an isopropanol precipitation. Seventy-five bp single-ended reads were generated on the Illumina NextSeq 500 platform at the Ramaciotti Centre for Genomics (UNSW, Sydney, Australlia).

### DNA extraction and sequencing for assembly

Genomic DNA was extracted from 13 mg of ethanol-preserved muscle tissue from a male (XY) *B. duperreyi*, using the Circulomics Nanobind tissue kit as per the manufacturer’s protocols, including the specified pre-treatment for ethanol removal. Library preparation was performed with 3 µg of DNA as input, using the SQK-LSK109 kit from Oxford Nanopore Technologies and sequenced across two promethION (FLO-PRO002) flow cells, with washes (EXP-WSH004) performed every 24 hours.

### Genome assembly

The genome assembly pipeline relied on using a combination of whole-genome sequencing ONT long reads, Illumina short reads and Hi-C reads. Firstly, adapters were removed from Illumina and Hi-C reads using TrimGalore v0.6.6 with default parameters. A primary assembly was generated using Flye v2.8.3 (34) and the ONT long reads, with the following parameters “--trestle --iterations 2”. Illumina short reads were aligned to the primary assembly with bwa-mem v0.7.17-r1188 and ONT long reads were aligned with minimap2 v2.24-r1122 (35), and both alignments were used to polish the assembly with hypo v0.5.1. Homologous contigs were identified and removed using Purge Haplotigs v1.1.1 (36). Chromosome-length scaffolding was performed with Hi-C data using Juicer v1.6 (37) to generate a contact matrix of the connections between contigs and 3d-dna v180922 to organise the contigs into larger scaffolds. Gaps in the assembly were filled using PBjelly v15.8.24 (38) with the ONT reads. The final assembly was assessed for completeness using BUSCO and accuracy using Merqury v1.3 (39), by comparing the assembly k-mer spectrum to those found in the Illumina reads.

### Annotation

The annotation used in this study was generated using AUGUSTUS v3.4.0 (40) with RNA sequencing data from brain, heart and gonad from male and female individuals. Prior to running AUGUSTUS, a transcriptome assembly and a soft-masked genome assembly were generated with Trinity v2.12.0 (41) and RepeatMasker v4.1.2-p1 (42), respectively. AUGUSTUS parameters were optimised with a training set of 500 genes, including 5 rounds of optimisation with exon and UTR parameters, before gene prediction and stitching. Peptides were inferred from the final annotation using gffread v0.12.7 (43) and blasted against the uniprot database with blast v2.11.0 (44) for gene identification. A github repository of this pipeline can be found at https://github.com/kango2/Annotation. Y specific sequences (20) in *B. duperreyi* were blasted against the generated genome assembly with blast v2.11.0 (44).

### Read depth analysis

DNA was extracted from muscle samples of the individual animals using the Gentra Puregene Tissue Kit (QIAGEN, Australia), following the manufacturer’s protocols with the modifications described below. The volume and reagent amounts were adjusted according to the size of the tissue sample, using three times more reagent than specified in the manufacturer’s protocols. Additionally, we made modifications to the DNA precipitation steps outlined in the manufacturer’s protocol. The DNA thread was spooled out using a galls rod and submerged in 300ul of 70% ethanol, then air dried for one minute. Subsequently, we used TE buffer for DNA hydration and allowed it to dissolve overnight at room temperature.

Raw read quality of the DNA sequencing was assessed using FastQC v0.11.9 (45) and trimmed accordingly with trimmomatic v0.38 (46). Trimmed reads were aligned to the newly generated *B. duperreyi* genome assembly with subread-align command in subread v2.0.1 (47). Read depth in 20000 bp windows was calculated using the bamCoverage command, from deeptools v3.5.1 (48), for the X and Y scaffolds as well as a representative autosomal scaffold (contig 328). The generated BED files were then used to plot read depth across these scaffolds using the ggplot2 package in R v4.2.1.

### Bioinformatics analysis of RNA-seq

Raw read quality of RNA sequencing was assessed using FastQC v0.11.9 (45) and trimmed accordingly with trimmomatic v0.38 (46). Trimmed reads were aligned to the newly generated *B. duperreyi* genome assembly with subread-align function in subread v2.0.1 (47). The subread-featurecount function from subread v2.0.1 (49) was used to count reads that overlapped genomic features using the settings “-0 -2 -t CDS,five_prime_utr,three_prime_utr”, the remaining settings were left as default. Library size and read counts were normalised by counts per million and RPKM respectively using the edgeR package in R v4.2.1 (50). Gene expression ratios were calculated and plotted with the ggplot2 package. Gene expression values were calculated from the RNA-seq data (hatchling brain tissue) of XX females (n=3), XY males (n=3) and sex reversed XX males (n=2). Normalised expression values were used to create male to female median expression ratios between the three genotypic and phenotypic sexes for each gene. Genes were binned according to their location on an autosome, the PAR, X as total output and X genes with Y gametologues (XY shared).

### Gene ontology and gene function

Gene ontology analysis was performed using g:Profiler (51) with gene lists identified using the *B. duperreyi* annotation. Results were restricted to biological process and molecular function and used the human database.

Protein functions of Y genes were obtained using the panther database v19.0 (52).

## DECLARATIONS

### Ethics approval and consent to participate

The animals involved in this study were collected from nests in the field under permits issued by the Australian state government: ACT permit numbers (LT201826, LT2017956) and NSW government (SL102002). The collection was carried out in accordance with their respective guidance and protocols, as well as the University of Canberra Animal Ethics Committee (Approval Number AEC 17–26). All husbandry practices complied with the Australian Code for the Care and Use of Animals for Scientific Purposes, 8th edition (2013), particularly sections 3.2.13–3.2.2.

### Consent for publication

Not applicable

### Availability of data and materials

All raw sequencing data generated in this study have been submitted to the NCBI BioProject database (https://www.ncbi.nlm.nih.gov/bioproject/) under accession number PRJNA980841. The genome assembly presented in this study is available from the NCBI Genome database under GenBank accession GCA_041722995.1.

### Authors’ contributions

B.J.H wrote the original draft. P.D.W and A.G designed the study. D.S.B.D collected and harvested samples. B.J.H and J.K.C conducted extraction experiments to generate data. A.L.M.R, J.M.H and J.K.C generated the genome assembly and annotation. B.J.H and J.K.C conducted data analysis with input from A.M.M and N.C.L. D.S.B.D, I.W.D, A.R-H, H.R.P, J.A.M.G and A.G provided insight into data interpretation and writing. All authors reviewed and approved the submitted manuscript.

## Acknowledgements

Sequencing costs for this project were met by grants from Bioplatforms Australia (AusARG initiative) and the Australian Research Council (DP170101147). We acknowledge the Garvan Sequencing Platform.

## SUPPLEMENTAL FIGURE LEGENDS

**Supplemental Figure 1** Blastn hits to the the largest 15 scaffolds representing the *B. duperreyi* chromosomes with males specific kmer contigs.

**Supplemental Figure 2.** Expression ratios for each autosomal scaffold for the three pairwise sex/ genotype comparisons presented as boxplots. Number of genes (n) is show in the key for each contig.

**Supplemental Figure 3.** Expression ratios were calculated for each gene on **A**) a representative autosome (contig 31) and **B)** the X chromosome (contig 1241), in a pairwise fashion for each sex (XYm, XXm and XXf). Values are plotted on a log2 scale and median values for each comparison are plotted as a red line. For panel B median for the X specific region is a red line and PAR is a blue line.

**Supplemental Figure 4** Scatterplot of counts per million (CPM) for each gene for each pairwise comparison of XXf to XYm, XXm to XYm and XXf to XXm. X genes are plotted in black with slope plotted as a red line. Autosomal genes are plotted in blue with slope as a green line. Slope and R-squared of each trendline is shown above each plot.

**Supplemental Table 1** List of genes on the Y chromosome (contig 93) including protein function

**Supplemental Table 2** Gene ontology results for XY shared genes

**Supplemental Table 3** Gene ontology results for all Y genes

