## Supplementary figures and images for "Sex chromosome dosage compensation in a sex reversing skink is not influenced by sexual phenotype"

### Supplemental Figure 1

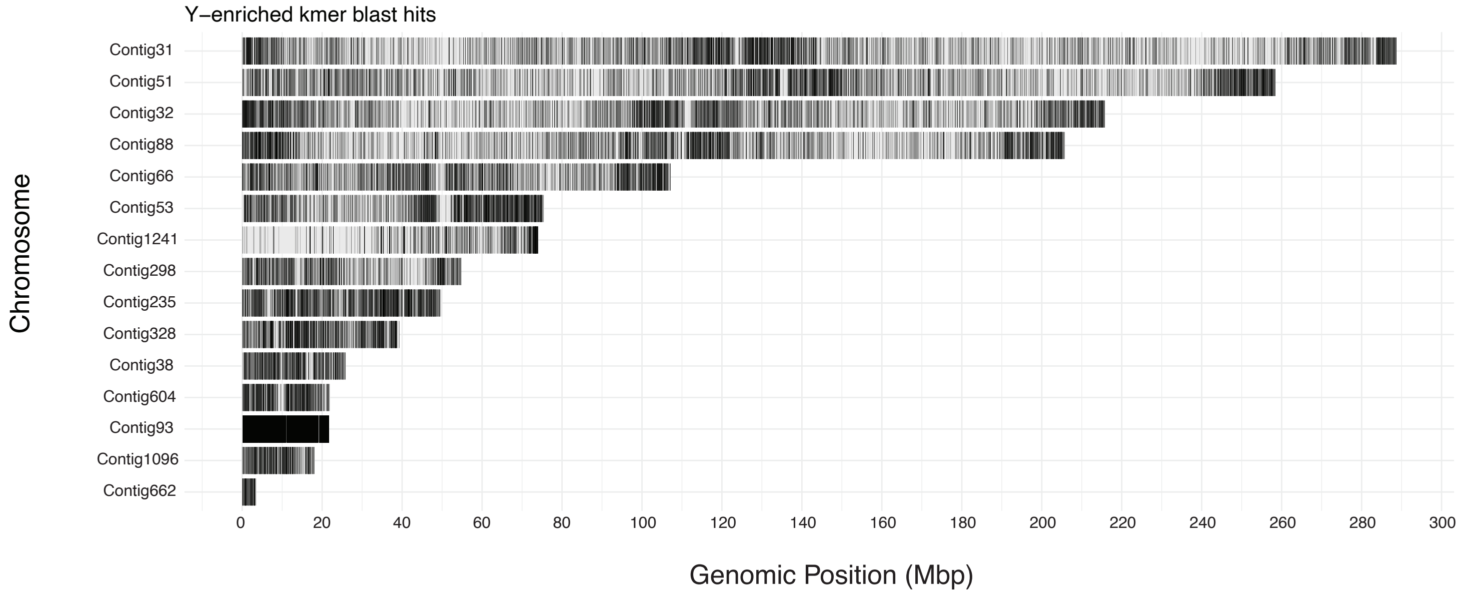

### Supplemental Figure 2

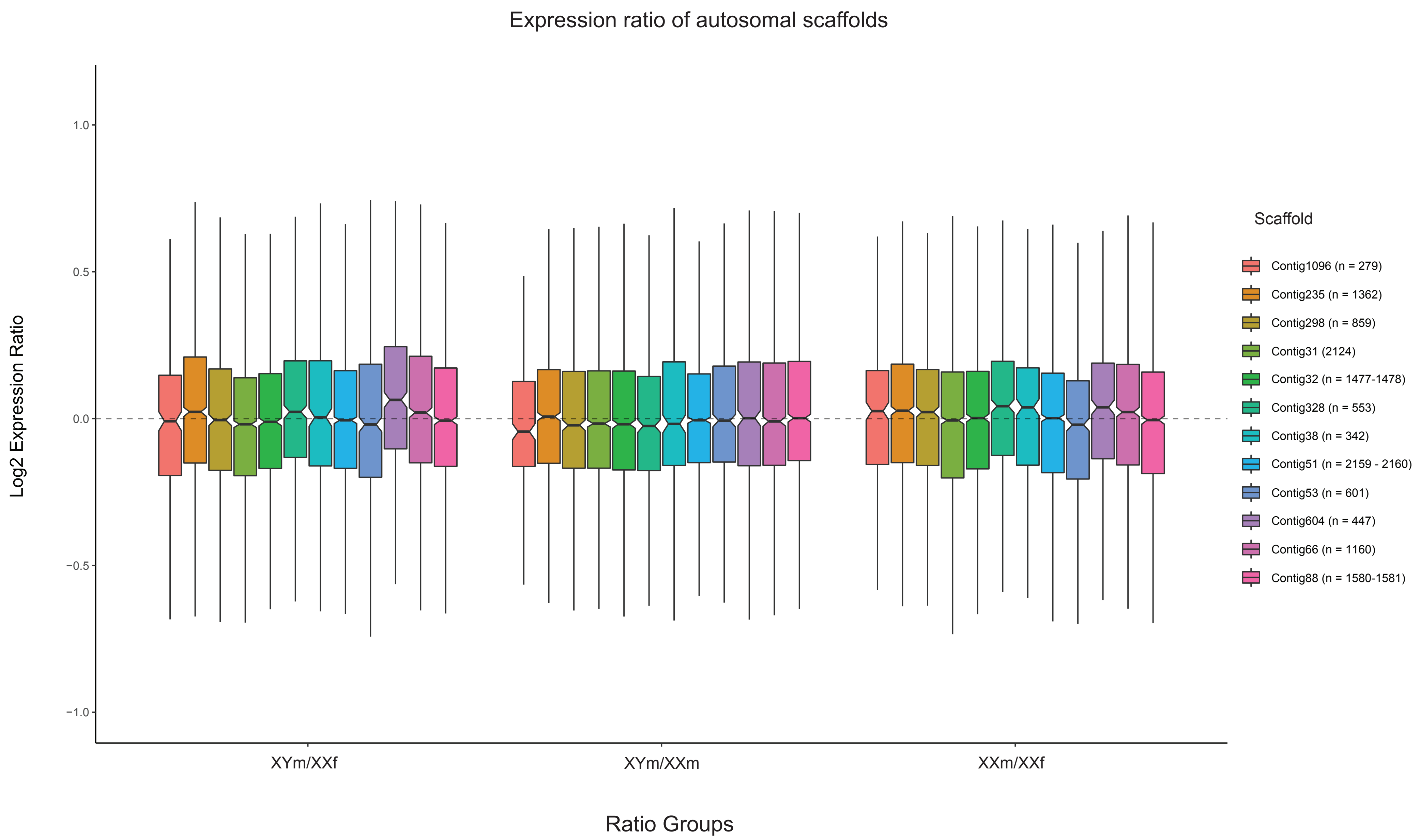

### Supplemental Figure 3

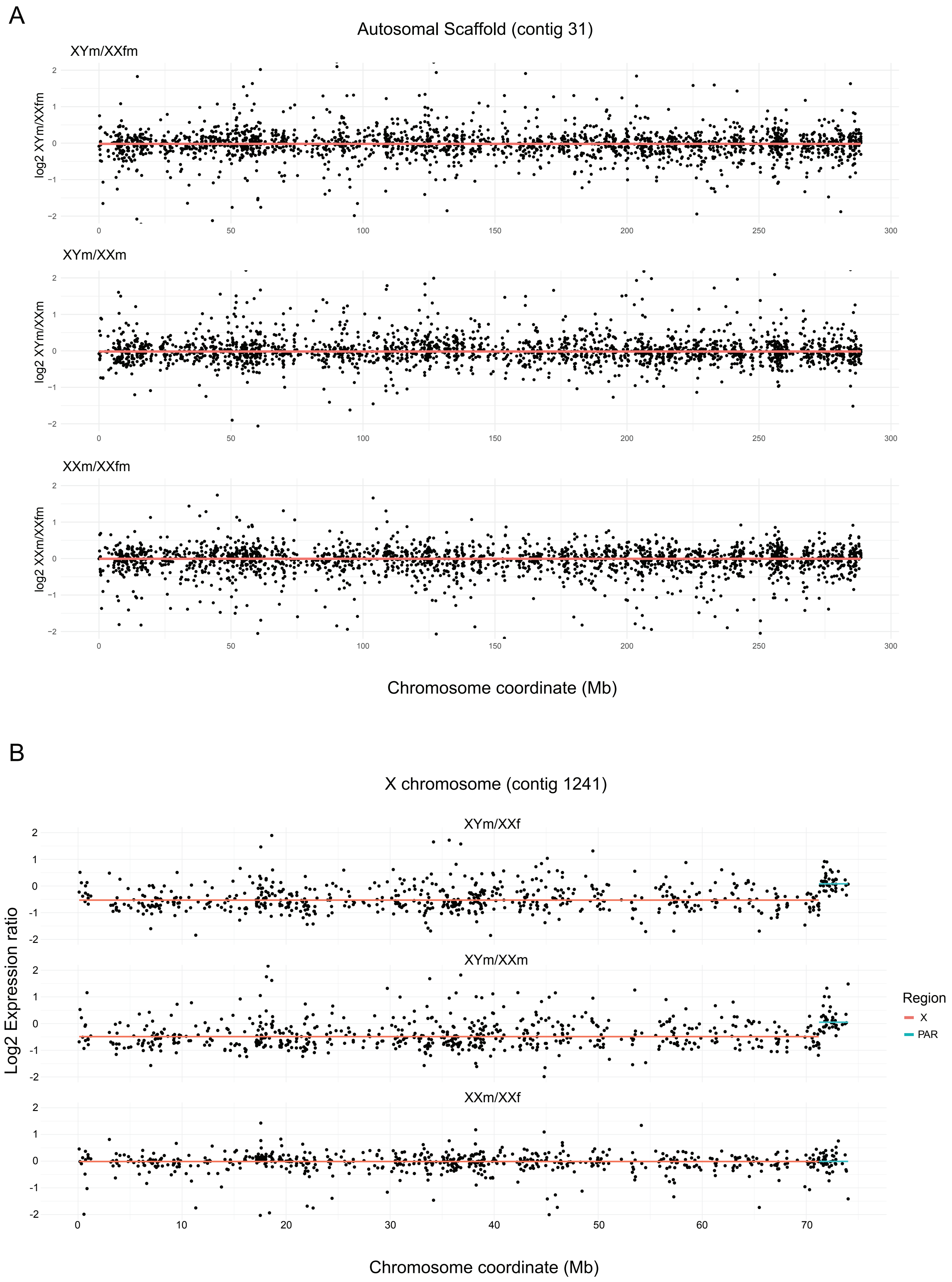

### Supplemental Figure 4

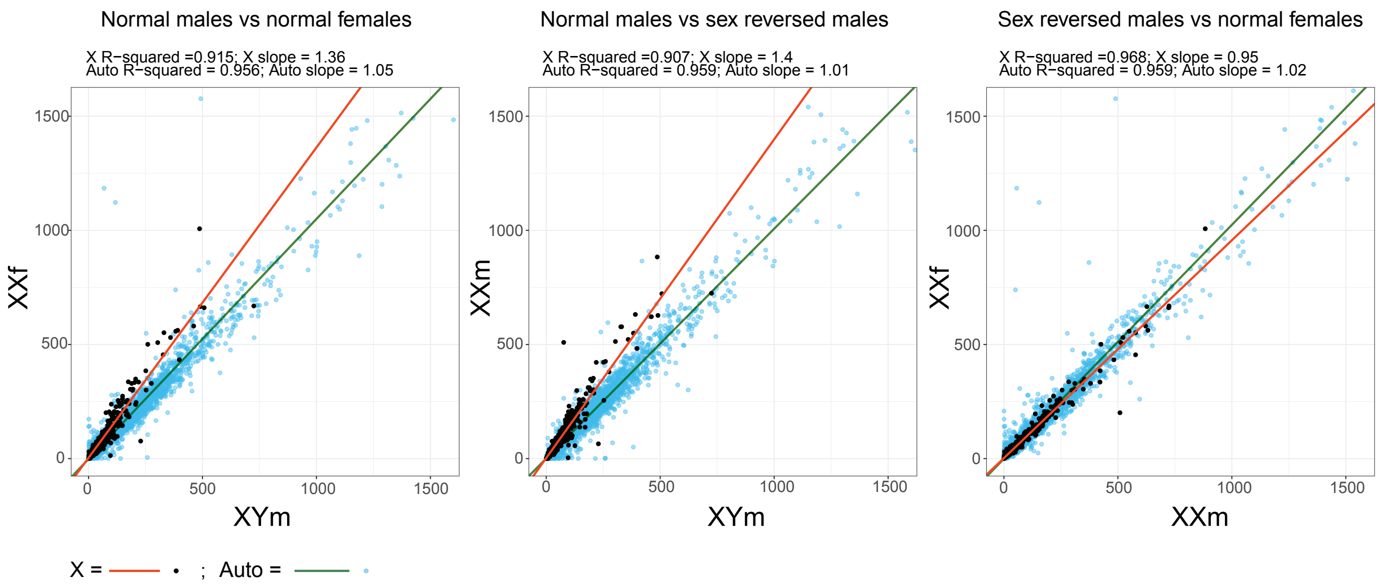
